# An electrochemical biosensor for rapid detection of anti-dsDNA antibodies in absolute scale

**DOI:** 10.1101/333641

**Authors:** Pablo Fagúndez, Gustavo Brañas, Justo Laíz, Juan Pablo Tosar

## Abstract

Autoimmune diseases are chronic inflammatory pathologies that are characterized by the presence of antibodies against own epitopes in serum (autoantibodies). Systemic lupus erythematosus (SLE) is a common autoimmune pathology, characterized by the presence of antinuclear antibodies (ANAs). These include anti-dsDNA (α-dsDNA) antibodies, which are widely used for diagnosis and disease monitoring. Their determination is carried out by traditional techniques such as Indirect Immunofluorescence (IFI) or Enzyme Linked Immunosorbent Assay (ELISA), which are time consuming, require qualified technicians, and are not compatible with decentralized analysis outside a laboratory facility. Here, we show a sandwich-format electrochemical biosensor-based method for α-dsDNA determination in a rapid and simple manner. Total assay time is only 30 minutes and the sensor is capable of detecting 16 ng (8 μg / mL) of α-dsDNA antibodies. Using the current derived from the detection limit of the method as a cut-off, we could discriminate positive from negative serum samples with 90% sensitivity and 100% specificity. By using monoclonal antibodies for calibration curves, our results are presented in absolute scale (i.e., concentration instead of serum title) what will help to perform comparisons between methods and further improvements of this protocol. In an effort to render the sensor compatible with automation, we minimized manipulation steps without compromise of the analytical performance, even in complex samples such as serum.

## Introduction

Autoimmune disorders are chronic inflammatory pathologies that affect over 5-8% of the world population^1^. In these disorders, the immune response of the individual is directed against its own components causing tissue- or organ-specific damage, generating local or systemic responses. Common autoimmune diseases include systemic lupus erythematosus (SLE), rheumatoid arthritis, multiple sclerosis, systemic sclerosis, type 1 diabetes, inflammatory bowel disease, and antiphospholipid syndrome. Diagnosis of these diseases is challenging for professionals, since their symptoms can vary between individuals and overlap with other rare pathologies^2,3^. Moreover, their origin is still now difficult to understand because of the genetic and environmental factors involved in their appearance ^4^. However, they share some common features, such as the presence of antibodies directed against own epitopes (autoantibodies).

SLE, considered as a model of autoimmune diseases, has been extremely studied and it has been reported more than a hundred autoantibodies which involve reactivity against nuclear, cytoplasmic and membrane components. The antinuclear antibodies (ANAs) are reactive against single and double strand DNA, histones, nucleosomes and chromatin, as well as other nuclear antigens (Ro, La and Sm ribonucleoproteins).^5^ Specific antibodies are associated with distinct clinical features. For example, the anti-dsDNA antibodies (α-dsDNA) are associated with the development of Lupus nephritis. Total ANAs, α-dsDNA and α-Sm antibodies are considered as the hallmarks of this pathology, and are included in the serological American College of Reumathology (ACR) criteria for diagnosis.^6–10^ SLE is also characterized by flares and remissions steps ^8,11^. Thus, the determination of the aforementioned antibodies in serum samples is relevant not only for diagnosis but also for classification, determination of the state of the disease, as well as for therapeutic evaluation and drug adjustment. ^3,12^

The gold standard technique for ANA determination is *Crithidia luciliae* indirect immunofluorescence (CLIF). In this assay, fluorescence intensity is used for title determination, and, in addition, it can be defined the specific autoantibodies contained in the ample because of the different staining patterns. Although several commercial kits have been developed with different cellular substrates, these require complex instrumentation and experienced stuff, being unsuitable for *point-of-care* applications.^13–15^ Recently, Enzyme Linked Immunosorbent Assay (ELISA) kits have been developed and commercialized. These kits use single or multiple antigen-coated wells (purified or synthetic), such as Ro, La, and dsDNA. However, diversity in antigen and adsorption strategies can affect comparability of the results between labs using kits from different suppliers. Line immuno semi-quantitative assays (variation of immuno-blots) are easy to use and facilitate detection of multiple autoantibodies in one strip. Nevertheless, they generally offer low sensitivity and specificity for certain antibodies.^16^

All the assays described above are time-consuming or can only be developed in a centralized laboratory with certificated equipment and technicians, being unable to provide immediate results for flare prediction and drug adjustment therapy. Biosensors emerge as suitable platforms for quick *point-of-care* tests. A biosensor can be described as an analytical device that includes a biological component as sensor, which works in association to a physicochemical transducer.^17,18^ Most biosensors oriented to the detection of autoantibodies employ a dsDNA-coated surface (or other auto-antigen) which is used to capture the autoantibodies in the sample. These can be detected directly (label-free) or indirectly (e.g., with the help of an enzyme-coupled secondary antibody). Some label-free approaches include Surface Plasmon Resonance chips (SPR) or Quartz Crystal Microbalance (QCM) measurements.^13,19–21^ However, electrochemical label-based biosensors are very promising for *point-of-care* tests because of the low costs associated with their automatization and miniaturization. ^22^ In this regard, Konstantinov, Rubin and coworkers have developed an electrochemical sandwich-type immunoassay where nuclear antibodies are detected through current measurements (using redox enzyme-coupled secondary antibodies). ^23–25^

Recently reported electrochemical biosensors as well as commercial kits use a WHO-standardized serum as a reference to measure autoantibody concentrations. Thus, concentrations are reported either in arbitrary units or in title units (maximum serum dilution to obtain a signal with a suitable signal-to-noise ratio), which are compared against the WHO reference or other previously calibrated positive samples. This is suitable for most clinical applications, but we argue that results expressed in absolute scale (i.e., molarity or grams per liter) are convenient for both basic research and applications where the inclusion of a previously calibrated control is not always possible (e.g., self-monitoring). Furthermore, the WHO-standardized reference serum is not available any longer.^2^

The previous reasons pushed us to develop an amperometric biosensor capable of detecting α-dsDNA antibodies, which could serve as a diagnostic and disease tracing tool in the near future. We also propose a methodology to obtain absolute α-dsDNA antibody concentrations (in terms of mass per volume), without the need of positive and previously calibrated human sera.

## Materials and methods

### Reagents and solutions

Lyophilized genomic salmon sperm DNA (“dsDNA”, Sigma-Aldrich, Cat.No. D1626) was employed for ELISA assays and electrode modification. Poly-L-Lysine and 3,3′,5,5′ tetramethylbenzidine (TMB, ≥98%) were purchased from Sigma-Aldrich. Freeze-dried bovine serum albumin (BSA ≥96%) was obtained from Spectrum Chemical Mfg Corp. (USA) and 1% fresh solutions were made in phosphate buffer saline (“PBS”: 10mM sodium phosphate buffer, 150 mM NaCl, pH = 7.4). Mouse monoclonal anti-dsDNA antibodies (α-dsDNA: sc-58749), normal mouse immunoglobulins (m IgG: sc-2025) and mouse monoclonal anti-human CD9 antibodies (α-CD9: sc-13118) were purchased from Santa Cruz Biotechnology INC. Rabbit HRP-conjugated anti-mouse IgG (α-m-IgG/HRP, ab6728) was purchased from Abcam. Fetal bovine serum (FBS) was supplied by Gibco. Stock DNA concentrations were estimated using a NanoDrop 1000 spectrophotometer (Thermo Scientific). Screen-printed three or four (bi-electrode) electrode strips (work: carbon; counter: carbon, pseudoreference: silver) were supplied by DropSense (Oviedo, Spain). All electrochemical measurements were carried out with a CHI760D workstation (CH Instruments).

### ELISA assay

The ELISA assay was carried out as previously described.^26^ A 96-well ELISA plate (CELLSTAR, Cat.-No.655 180) was pretreated with 100 μL of 50 μg/mL poly-L-Lysine solution (in water) for 30 min at room temperature (RT) and washed three times with Tris Borate Saline buffer (“TBS”, 10 mM, 150 mM NaCl pH = 7.5). Then, 100 μL of dsDNA solution (4 μg/mL in TBS) was incubated for 60 min at RT and washed in the same way as in the previous step. After this, the plate was blocked over-night with 100 μL of 1% BSA solution at RT, and washed vigorously three times with PBS-0.1% Tween-20 buffer. Then, 100 μ L of PBS dilution containing either α-dsDNA or m IgG (specific and unspecific antibodies respectively) was incubated for 60 min at 37ºC and washed with PBS - 0.1% Tween as previously described. 100 μL of the α-m-IgG/HRP (1/2000 dilution, plus 1% BSA) was dispensed and incubated for 60 min at 37ºC. The plate was washed and finally incubated for 30 min with 100 μ L of TMB-H_2_O_2_ solution (2 mM TMB, 1mM H_2_O_2_, diluted in 50 mM acetic acid/sodium acetate buffer, pH = 5). The reaction was stopped with 50 μL of 5M HCl and the absorbance at 450 nm was measured in a microplate reader (Thermo Scientific, Multiskan EX).

### Construction of dsDNA-modified electrodes

100 uL of acetic acid/sodium acetate buffer (200 mM, pH = 5) was placed in order to cover the three or four screen-printed electrodes in each strip, and a potential of +1.7 V was applied for 120 s as electrode pretreatment, followed by three washes with the same buffer. A dsDNA solution (0.5 μg/μL in acetic/acetate buffer) was prepared and vortexed for 30s and then 50 μ L were placed on the electrode system. A constant potential of +0.5 V was applied for 300 s for dsDNA immobilization. After washing three times with PBS, 2 μ L of 1 % BSA were deposited (on the working electrode only) and incubated for 30 min at 37 ºC in a wet chamber with 100% humidity.

The dsDNA immobilization was confirmed by cyclic voltammetry (CV) after successive washing steps. Briefly, 50 μ L of acetic acid buffer were placed on the three electrode system and CV was carried out between −0.2V and +1.0V at a scan rate of 0.05 V/s in order to observe the guanine oxidation signal.

### Detection of specific antibodies

Different dilutions of α-dsDNA and m IgG were prepared in PBS. Two microliters of these solutions were placed on the working electrode and incubated for 20 min at 37ºC in a wet chamber. For the bi-electrode strips, each antibody was placed in a different working electrode. After this, the electrodes were washed three times with PBS and 2 μ L of α-m-IgG/HRP antibody (1/2000, 1%BSA) was added and incubated under the same experimental conditions. The electrodes were then washed with PBS and 50 μ L of a TMB/H_2_O_2_ solution was added. Immediately, the working electrode was placed at a constant potential of −0.1V and the TMB reduction current was registered. For optic measurements, only monoelectrodes were employed. After antibody incubations, the TMB/H_2_O_2_ solution was incubated for 5 min, followed by the addition of 5 μ L of 5M HCl. Two microliters of the mixture were measured at 450 nm using a NanoDrop spectrophotometer.

### Performance in serum samples

As an approximation to the analysis of real samples, assays were performed in the bioelectrode strips by spiking α-dsDNA or irrelevant antibodies directly in 1/80 FBS. Here, two different strategies were tested. One strategy (which we called the “two-step” method) consisted in incubating α-dsDNA-containing FBS on the working electrode, performing two wash steps with PBS, and then incubating (20 min at 37°C) the electrodes with 2 μ L of α-m-IgG/HRP antibody (1/2000 dilution, plus 1% BSA) followed by a new washing step and addition of the TMB/H_2_O_2_ solution. In contrast, the “one-step” method consisted in preincubating the α-dsDNA (or irrelevant antibodies) with the α-m-IgG/HRP in FBS. Briefly, 5 μ L of α-dsDNA (in 1/40 FBS) was mixed with 5 μ L of α-m-IgG/HRP (1/1000) and incubated for 5 min at 37ºC. Then, 2 μ L of the mixture was placed on the working electrode and the procedure continued the same way as described above. As a consequence, one single incubation with the sample and HRP-conjugated antibodies was needed, avoiding the washing step in between, and reducing assay timeapproximately two-fold.

## Results

We propose the construction of an amperometric biosensor capable of detecting anti-double stranded DNA antibodies, which offer diagnostic and prognostic value in several auto-immune diseases. This biosensor is based on the specific binding of α-dsDNA antibodies present in a test sample to dsDNA molecules immobilized on the surface of a disposable screen-printed carbon graphite electrode, and the subsequent binding of anti-mouse IgG antibodies conjugated to the electroactive enzyme HRP (conjugated ABs). The electrochemical reduction the oxidized TMB generated by the catalysis of HRP is measured^27^, what is related to the amount of α-dsDNA antibodies located in the sensor’s surface. In an attempt to render this biosensor compatible with automatization, the conjugated ABs were directly introduced in the test sample, greatly simplifying the detection procedure (one-step method, **Scheme 1**).

### Specificity of the α-dsDNA antibody

Specificity of our α-dsDNA antibody was studied by ELISA (**Figure 1**). Wells sensitized with either vortexed or intact dsDNA showed a characteristic binding response when incubated with increasing concentrations of the mouse monoclonal α-dsDNA antibody, with a linear response below 0.07 μg / mL. In contrast, incubation with normal mouse immunoglobulins (m IgG) showed basal absorbance independently of the concentration used, confirming specificity of the assay. Secondly, we assayed the α-dsDNA antibody against genomic DNA, plasmid DNA and a purified PCR product, confirming that it is the dsDNA molecule itself (rather than co-purified nucleosomes present in certain genomic DNA preparations) which is being recognized by the antibody used throughout this study. (**Supplementary Figure 1**).

**Figure 1.**
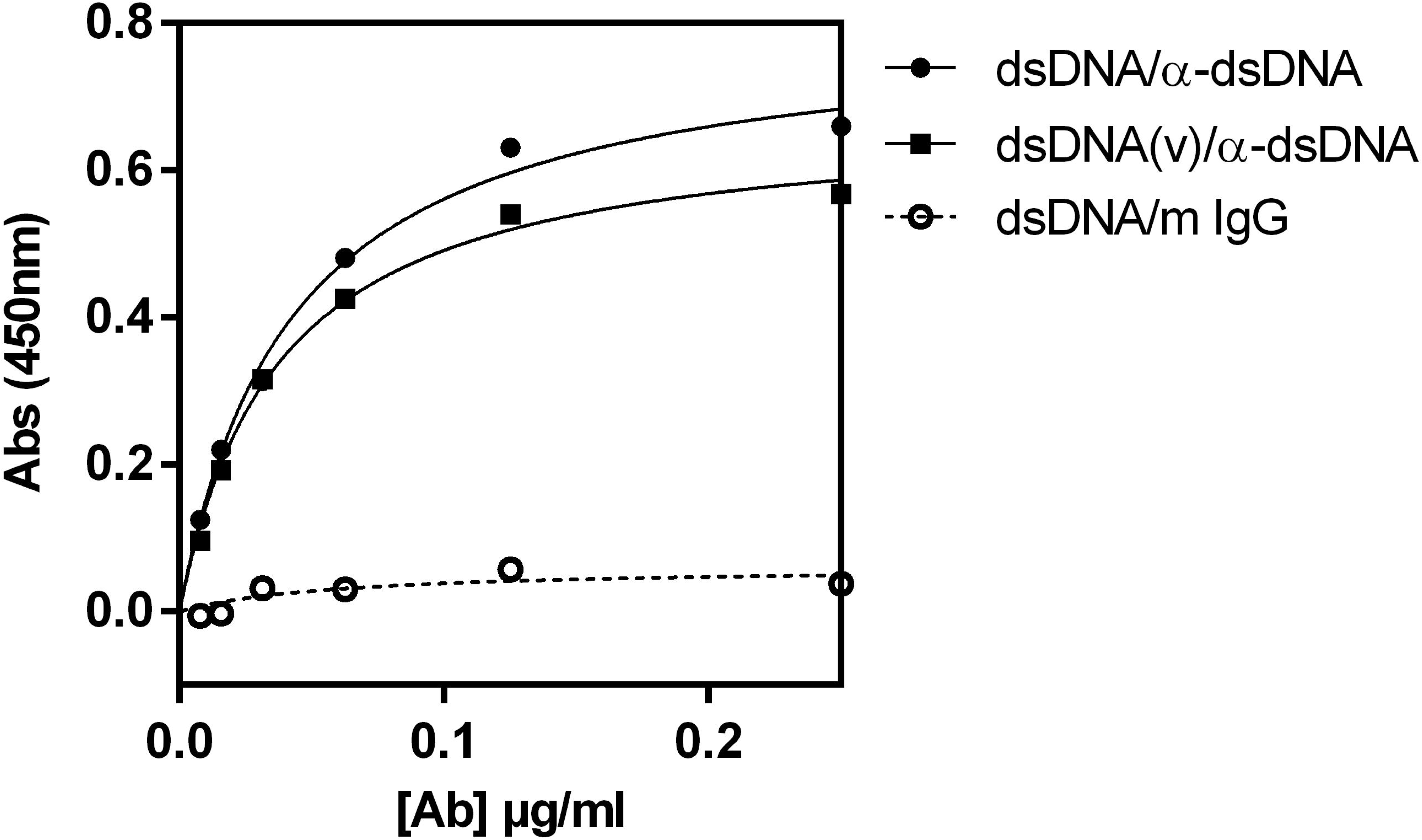
Specificity of α-dsDNA antibody by enzyme-linked immunosorbent assay. Plates were sensitized with vortexed (squares) or intact (circles) salmon sperm DNA and incubated with different concentrations of α-dsDNA antibody (black) or mouse IgGs as a control (white).

### Electrode modification

Deposition of dsDNA on the working electrode was done at a constant potential as indicated in Methods. We asked whether that immobilization method would be suitable for maintaining the dsDNA attached to the electrode surface during the whole procedure. To test this, we subjected dsDNA-modified BSA-blocked electrodes to a number of washes in PBS, which was equivalent to the number of washes in our longest assay (three washing cycles, three washes per cycle). The presence of DNA in the electrodes was then analyzed by measuring the irreversible guanine oxidation signal^28^ at +0.83 V by cyclic voltammetry, and comparing this signal with naked (**Figure 2A**) or BSA-only electrodes (data not shown).

**Figure 2.**
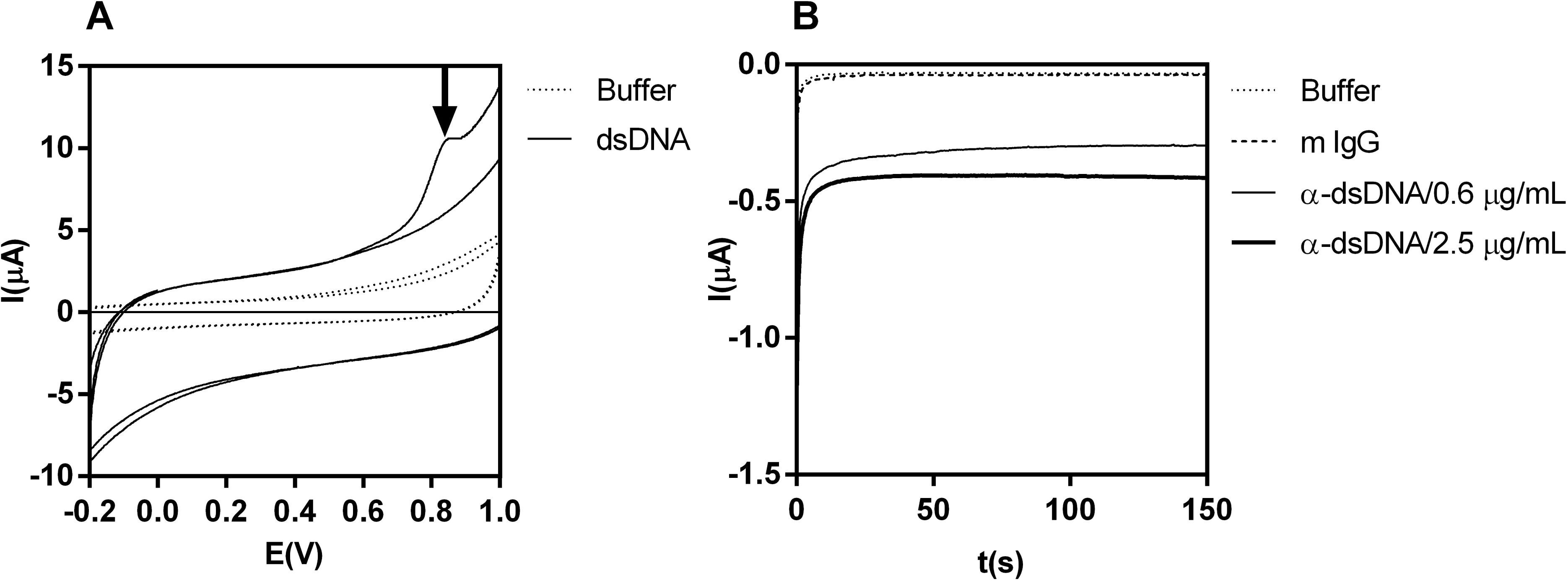
Electrode modification with dsDNA and electrochemical sensing of α-dsDNA antibodies. A) cyclic voltammograms showing guanine oxidation (arrow) in dsDNA-modifiedelectrodes (solid line) vs buffer-treated (dashed line) screen-printed carbon electrodes. B)Intensity vs time curves for TMB reduction on the electrode surface at – 0.1 V. dsDNA-modified BSA-blocked electrodes were incubated with α-dsDNA antibodies at two different concentrations (solid lines), normal mouse IgG at 2.5 μg / mL (thick dashed line), or buffer (thindashed line), and then incubated with α-mouse IgG/HRP conjugates, followed by addition of a TMB/H_2_O_2_ solution.

### Specificity of the electrochemical biosensor

The electrochemical behavior of TMB on our screen-printed dsDNA-modified carbon electrodes was studied by cyclic voltammetry (**Supplementary Figure 2**) in order to determine the working potential where electrochemical TMB oxidation was virtually zero, and also registering the electrochemical reduction of oxidized TMB. By doing so, we defined −0.1 V as a suitable applied potential for subsequent constant potential assays.

As a proof-of-principle, intensity vs time (I vs t) assays at −0.1 V are shown in **Figure 2B**. Incubation of dsDNA-modified electrodes with the α-dsDNA monoclonal antibody (followed by detector ABs and TMB/H_2_O_2_ solution) showed the expected response ^27^, with a negative (i.e., reduction) signal decreasing in absolute value with time, and approaching a concentration-dependent plateau as predicted by the Cottrell equation.^29^ Here, total TMB concentration is constant, but the concentration of oxidized TMB depends on the number of HRP molecules present at the electrode’s surface. In contrast, electrodes incubated with normal mouse IgG displayed a negligible signal, comparable to electrodes only exposed to the conjugated ABs. This finding demonstrates that unspecific binding of conjugated ABs to our electrodes is despicable in the absence of analyte (i.e., α-dsDNA antibodies).

### Optical and electrochemical response

Based on the fact that our electrochemical signal depends on the concentration of enzyme-oxidized TMB, which also absorbs visible light with a maximum at 450 nm after acidification of the medium, we compared the analytical performance of the sensor by both optical and electrochemical readouts. Similarly to what was previously observed in ELISA plates (**Figure 1**), the absorbance of oxidized TMB was saturated above 0.05 μg / mL of α-dsDNA antibodies in carbon/dsDNA/BSA electrodes (**Figure 3A**), with no signs of unspecific binding of antibodies to the electrode surface. In contrast, the electrochemical response (measured as the electrical current in I vs t plots at exactly t = 100 s) was linear up to approximately 0.5 μg / mL, showing a 10-fold increase in the dynamic range of the method when compared to absorbance measurements. It is important to mention that this assays where performed in bi-electrodes, where both working electrodes were modified simultaneously using the same dsDNA solution, and incubated with either α-dsDNA antibodies or normal mouse IgGs at exactly the same concentration. Thus, batch effects (i.e., disparities in electrode fabrication and dsDNA immobilization) are minimized due to the use of paired data in our assays.

**Figure 3.**
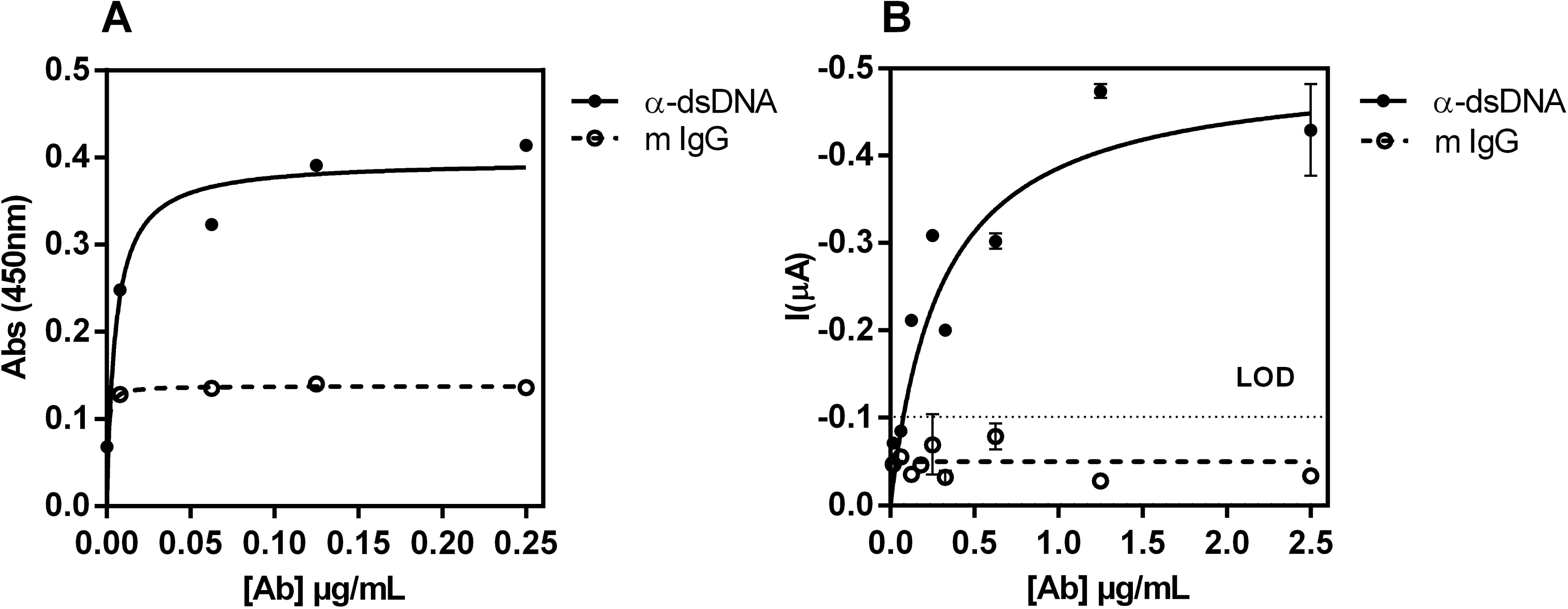
Optical and electrochemical response of the sensor. A) Absorbance changes as a function of the concentration of m α-dsDNA (black circles) or mouse IgGs (open circles). B) Reduction current of TMB at 100s as function of the concentration of α-dsDNA (black circles) or mouse IgGs (open circles). LOD (limit of detection) was determined as the background signal plus three standard deviations. For visualization purposes, the current axis direction was inverted. Error bars correspond to the standard error of the mean of two independent replicates. Bi-electrodes containing specific and irrelevant antibodies at the same concentration were assayed in parallel.

By using the standard deviation of all electrodes incubated with mouse IgGs, we determined the limit of detection (LOD) of our sensor as the background signal (i.e., the average of mouse IgG-incubated electrodes) plus three standard deviations. We obtained a LOD of 0.1 μg / mL of α-dsDNA antibodies, corresponding to a measured current of - 0.1 μ A. Currents (cathodic) higher than that could be taken into account to discriminate samples that contain α-dsDNA antibodies.

### Performance in serum samples

Once addressing the sensitivity and specificity of the sensor towards α-dsDNA antibody detection in buffer, we wondered whether this device would be able to detect these antibodies in a complex sample mixture such as serum. It is reported that many sensors fail to give a well-response signal when exposed to real complex matrixes even if their performance indicates the recognition of specific analytes when using pure laboratory samples.^30^ To test this, we performed standard additions of the monoclonal α-dsDNA antibody (or irrelevant antibodies) in 1/80 dilutions of fetal bovine serum. We decided to work in 1/80 serum as this is the dilution used to differentiate positive from negative samples in SLE ^14,15^. Even though we could no longer detect a clear trend in the electrical current as a function of the specific antibody concentration, we did observe a statistically significant (*p < 0.0001*) higher reduction (i.e., negative) current of TMB in electrodes incubated with α-dsDNA-containing FBS vs. mouse IgG-containing FBS (**Figure 4A**). This also depended on the presence of immobilized dsDNA on the surface of the electrodes, as electrodes modified with BSA alone showed background currents even when incubated with α-dsDNA-containing FBS (**Figure 4A, diamonds**). Assuming a cutoff value of 0.1 μ A (which corresponds to the measured current at the LOD), the biosensor was 100% specific (no false positives) and 82% sensitive (3 false negatives out of 17). Similar results were obtained for current measurements at 50 s and for quantization of the electrical charge passed through the electrode during 100s (**Supplementary Figure 3**)

**Figure 4.**
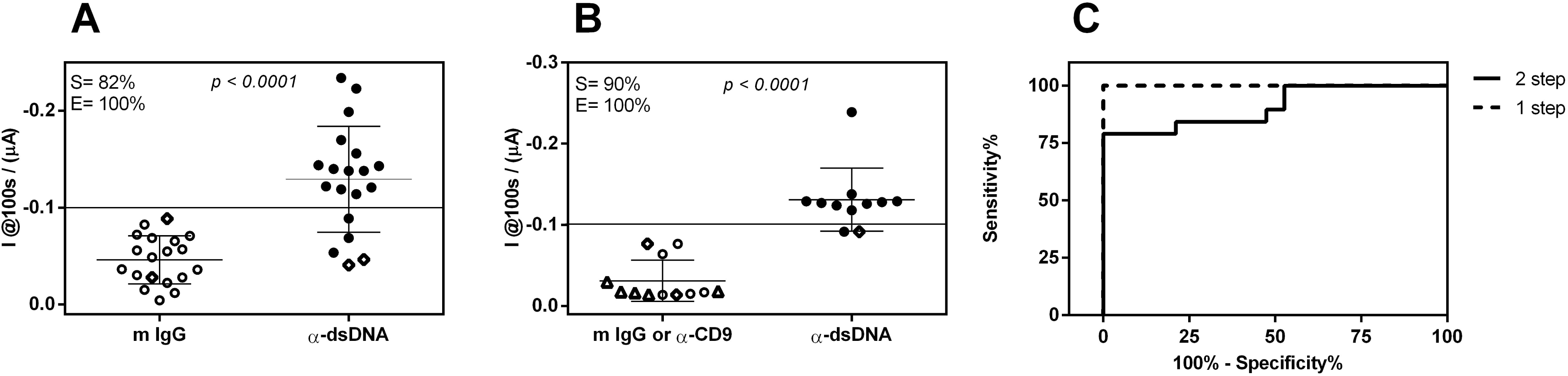
“One-step” vs “two-step” method. Reduction current recorder at 100s after incubation with serum samples containing either relevant or irrelevant antibodies at the same concentration with the “two-step” (A) or “one-step” (B) method. To reduce batch effects, specific (black circles) and irrelevant (mouse IgG: open circles; monoclonal α-human CD9, open triangles) antibody-containing samples were incubated in parallel in bi-electrode strips. Diamonds correspond to carbon electrodes not modified with dsDNA (negative controls). Error bars correspond to the standard error of the mean. Student *t* test (two-tailed) was carried out to test statistically significance in the difference between positives and negatives. The current corresponding to the detection limit of the method was used as cut-off in order to establish assay specificity and sensitivity. Alternatively, a mobile cut-off was used in order to obtain ROC (receiving operating characteristic) curves for both methods (C).

Our next attempt was to reduce assay time and manipulation steps, in order to ease automatization of the sensor in the future. We hypothesized that by adding the conjugated ABs directly into α-dsDNA-containing FBS, we could then incubate both molecules above the electrode surface in a single step, without the need for a washing step in the middle. This method, called “one-step” in opposition to the “two-step” method, reduced the total assay time to 30 min (vs. 60 min in the “two-step”) without compromising specificity (**Figure 4B**), as evident from a comparison of ROC curves for both methods (**Figure 4C**). In contrast, the current separation between positive and negative samples was even higher, with a 100% specificity and 90% sensitivity when using the 0.1 μ A cutoff. To discard that the positives results were not simply a consequence of using monoclonal antibodies instead of normal mouse IgG, we also analyzed other negative samples in which irrelevant monoclonal antibodies (e.g., anti-human CD9) were added to 1/80 FBS at the same concentration as the positive samples (**Figure 4B,** triangles). Overall, these results show that our biosensor is capable of discriminating between serum samples containing α-dsDNA antibodies and those which do not. This is promising towards the future development of electrochemical biosensors for autoimmune disease-based applications.

## Discussion

There is a need for rapid, simple and low-cost analytical devices capable of determining the presence and concentration of autoantibodies in blood specimens, to favor disease diagnosis, treatment and monitoring.^12,31–33^ Currently, these assays are carried out in clinical laboratories and demand time, costly equipment and specialized human resources. In contrast, electrochemical biosensors are intrinsically compatible with point-of-care testing^22,27,34^, and that makes them suitable for decentralized autoantibody determination. However, the number of reports describing electrochemical biosensors for anti-dsDNA autoantibodies (a hallmark of SLE and other autoimmune diseases) is rather scarce (**Table 1**).

**Table 1.** Assay time and limit of detection of different biosensors for analysis of autoantibodies.

| Biosensor/Transducer | Target | Time assay (min) | Sample amount* (μL) | LOD | Ref |
| --- | --- | --- | --- | --- | --- |
| QCM | α-TRIM21(Ro52)/α-TROVE2(Ro60) | > 60 | 1000 | NR | 20 |
| | α-dsDNA | NR | 2000 | NR | 21 |
| SPR | α-dsDNA | 15 | 45 | NR | 19;13 |
| Electrochemical | α-chromatin | 30 | 200 | NR | 25 |
|  | ANA | 30 | 200 | NR | 24 |
| | α-dsDNA | 30 | 200 | 10 μg/mL | 23 |
| | α-dsDNA (two-step) | 60 | 2 | 8 μg/mL | This work |
| | α-dsDNA (one-step) | 30 | | | |

QCM: Quartz Crystal Microbalance. SPR: Surface Plasmon Resonance
ANA: Total Antinuclear Antibody.
NR: not reported.
\*Diluted serum samples

Some of the important parameters to study when describing new analytical technologies or methods are the sensitivity (often expressed as the limit of detection) and specificity of the assays, as well as the performance of the method in real samples. Previous reports, such as the seminal work performed by Rubin and Konstantinov, have used sandwich immunoassays in fluidic devices to detect anti-dsDNA, anti-nuclear or anti-chromatin autoantibodies with electrochemical biosensors based on a similar electrochemical readout as the used herein.^23–25^ The authors compared the amount of autoantibodies in patient serum samples, and stablished the performance of their sensor against commercial ELISA kits. However, their output was based on the value of the measured electrochemical current, so establishing comparisons of analytical performances with other methods was difficult, if not impossible. To overcome this problem, we performed calibration curves in artificial samples containing monoclonal anti-dsDNA antibodies, as suggested by Buhl et al. (2009).^2^ We obtained a limit of detection [LOD] of 0.1 μg of α-dsDNA antibodies per mL, and used the current in the LOD to define a cut-off in non-human serum samples were known quantities of specific or irrelevant antibodies were spiked-in. Through this procedure, we obtained a 90% sensitivity and a 100% specificity with the “onse-step” method (i.e., we could label 9 out of 10 positive samples as positive, and 10 out of 10 negative samples as negative). Considering the fact that we used 1/80 dilutions of serum, our real detection limit increases up to 8 μg/mL, which is slightly lower but comparable to previous reports^23^

Since reported detection limits are usually based on the title of autoantibodies (i.e., 1 / maximum dilution of a serum sample for which a positive signal is still detected) we wondered whether our detection limit of 8 μg / mL (16 ng total anti-dsDNA antibody, since sample volume is only 2 μ L) was clinically relevant. To study this, we performed serial dilutions of FBS spiked-in with known amounts of α-dsDNA antibodies, and looked for the amount which mimicked serial dilutions of a SLE patient serum by ELISA. By doing so, we estimated the α-dsDNA cargo in this patient sample to be precisely 8.8 μg / mL. Thus, the limit of detection of our sensor is in principle compatible with clinical applications. Further optimization will be needed to achieve this goal, as mammalian non-human antibodies and serum were used in this study.

One of the important aspects of our study is the minimization of assay time and manipulation steps achieved with the thus called “one-step” method. If electrochemical biosensors are chosen for their compatibility with point-of-care tests, they should be compatible with automatization in order to avoid the need of specialized technicians. By decreasing the number of manipulation steps (including sample incubation, washes, addition of reactants, among others) we facilitate the implementation of our sensing methodology in future analytical devices. By pre-incubating serum samples with the HRP-conjugated secondary antibodies, we have successfully decreased manipulation steps from 5 (sample incubation, wash, conjugate incubation, wash, TMB /H_2_O_2_ addition and electrochemical measurement) to 3 (sample incubation, wash, addition of redox mediator and electrochemical measurement). Furthermore, this protocol reduced total assay time from 60 to 30 minutes.

## Conclusion

We provide a simple protocol for the electrochemical sensing of α-dsDNA autoantibodies in serum samples, with excellent distinction between samples containing or not α-dsDNA antibodies in concentrations comparable to those present in the sera of autoimmune disease patients. The total assay time of 30 minutes and the few manipulation steps will aid in the automatization of this protocol in order to obtain portable sensors for their use outside of laboratory facilities.

## Acknowledgments

The authors want to thank Eduardo Méndez (Faculty of Science, Uruguay) for helpful discussions and Alfonso Cayota (Institut Pasteur de Montevideo) for some of the antibodies used in this work. JPT is a member of the National System of Researchers (ANII, Uruguay) and the Program for the Development of Basic Science (PEDECIBA, Uruguay).

## Figure Legends

**Scheme 1. Detection strategy.** Positive or negative serum samples (containing 1/80 fetal bovine serum plus either murine monoclonal anti-dsDNA [red] or irrelevant [green] antibodies) were incubated with dsDNA-modified carbon electrodes, washed, and later incubated with anti-mouse IgG antibodies conjugated to HRP enzyme [blue] (A). In a more time-efficient and automatization-compatible strategy, conjugated antibodies were mixed with the serum samples before incubation (one-step, B).

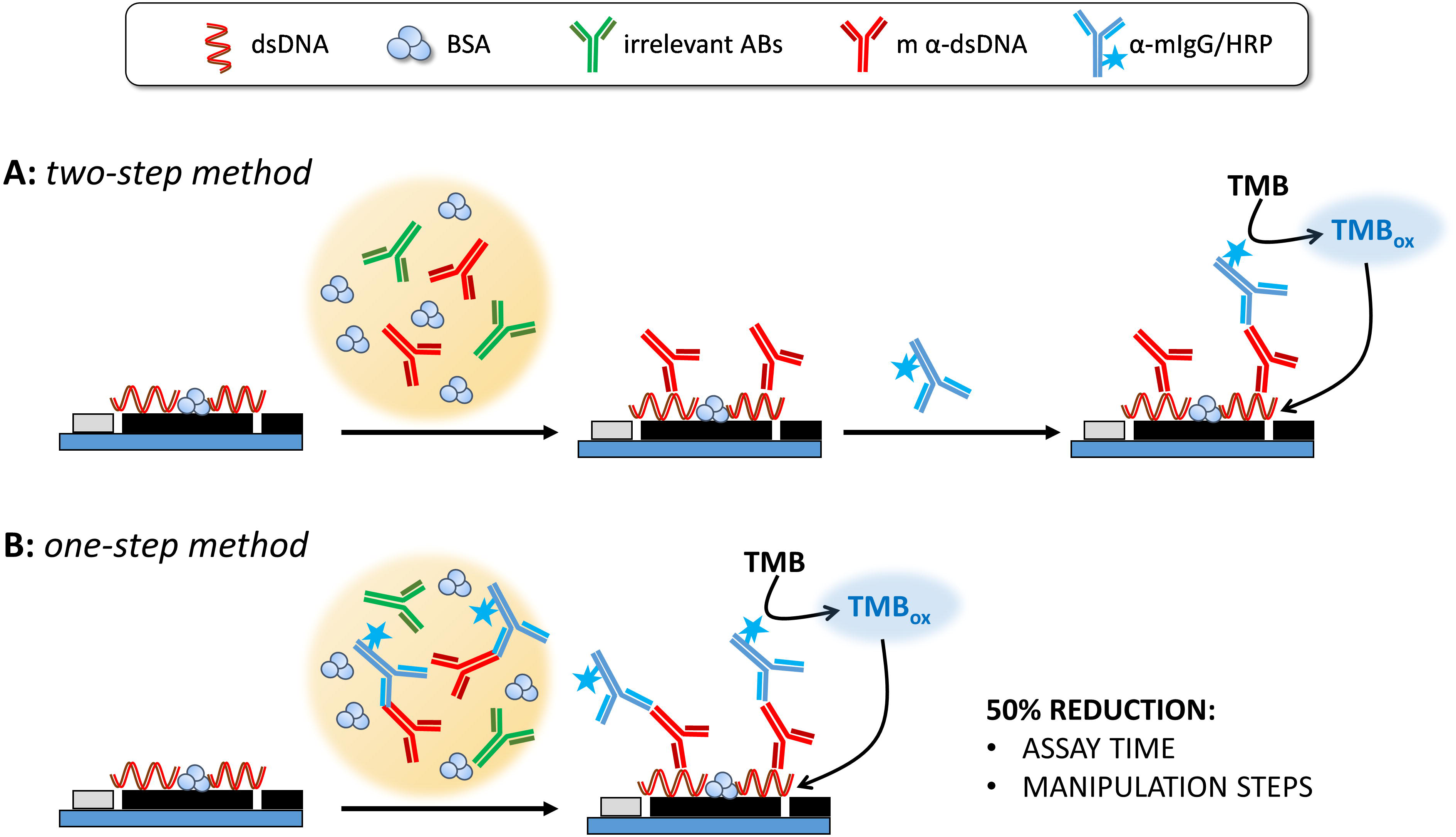

